# Long-read bitter gourd (*Momordica charantia*) genome and the genomic architecture of domestication

**DOI:** 10.1101/822411

**Authors:** Hideo Matsumura, Min-Chien Hsiao, Atsushi Toyoda, Naoki Taniai, Kazuhiko Tarora, Naoya Urasaki, Shashi S. Anand, Narinder P. S. Dhillon, Roland Schafleitner, Cheng-Ruei Lee

## Abstract

Bitter gourd (*Momordica charantia*) is a vegetable and medicinal plant of the family Cucurbitaceae. Here we report a chromosome-level assembly, with highest contig N50 (close to 10 Mb) and proportion of sequences placed on chromosomes (96%) in Cucurbitaceae. Population resequencing revealed the divergence between wild and cultivars at about 6000 years ago. Different cultivar groups have distinct allelic compositions in loci associated with domestication traits, suggesting phenotypic changes were achieved by allele frequency shifts in independent loci. Noticeably, one candidate locus for fruit size locates within a region missing from a recent Illumina-based assembly. Despite breeding efforts to increase female flower proportion, the gynoecy locus exhibits high variation within and low differentiation between wild and cultivar groups, likely because artificial directional selection could not overwhelm natural balancing selection. Our study provides resources to further investigate the genetic architecture of bitter gourd as well highlights the importance of a well-assembled genome.

## Introduction

Domestication involves human actively modifying organismal traits and is considered a good model to study the process of evolution (Meyer and Purugganan 2013). Classic examples include the *TEOSINTE BRANCHED 1* (*TB1*) gene generating non-branching of maize (Wang *et al.* 1999), the *QTL of seed shattering in chromosome 1* (*qSH1*) gene for non-shattering in rice (Konishi *et al.* 2006), as well as many others. Intriguingly, these “classic examples” involve strong directional selection on novel mutations of Mendelian traits, which left strong signatures of hard selective sweep. On the other hand, in most plants, domestication inevitably involves the enlargement of seeds or fruits, likely a highly polygenic traits where selection may only slightly altered the allele frequencies of standing variations. In some plants, the domesticated forms, semi-wild forms, and wild progenitors were all utilized by humans, and the continuum of phenotypic divergence is not as discrete as classic examples such as maize. The situation may be further complicated by the parallel selection in different countries, resulting in different sets of “domestication genes” for the same phenotype in cultivars of diverse genetic background. Therefore, to understand the process of domestication and how human might have shaped the genomes of plants, studies on these “non-classic” cases are necessary. Here we focus on bitter gourd (*Momordica charantia*, 2n = 2x = 22 (Zaman and Alam 2009)).

Bitter gourd is a vegetable and medicinal plant of the family Cucurbitaceae, cultivated in tropical and subtropical Asia and characterized by its spiny skin pattern and bitter taste. Bitter gourd fruits are rich in vitamin C, minerals, and carotenes (Urasaki *et al.* 2017). The pharmacological effect of bitter gourd has been widely investigated (Tan *et al.* 2016), especially in type 2 diabetes. Bitter gourd fruits contain substances with the antidiabetic effect such as charatin, vicine, and polypeptide-p, which may improve insulin sensitivity and decrease blood glucose level (Krawinkel and Keding 2006). While a good reference genome is strongly needed, the chromosome-level gnome of *M. charantia* based on long-read sequencing is not yet available. The most recent publicly available *M. charantia* genome is a short-read scaffold-level assembly (Urasaki *et al.* 2017) as well as another short-read based assembly connected by linkage map (European nucleotide archive PRJEB24032).

Previous studies have investigated the patterns of genetic variation of *M. charantia*: Five clusters were identified in the collection of India cultivars (Gaikwad *et al.* 2008), and three clusters were found using accessions from east and southeast Asia and 160 SSR markers (Saxena *et al.* 2015). The most recent study, using 50 SSR markers and 114 accessions, identified three major subgroups: India, Philippines, and Thailand (Dhillon *et al.* 2016). However, all population genetics study available to date used low-density markers. To fully investigate the demographic history of domestication as well as identifying important genomic regions under domestication and artificial selection, a population genomics study from re-sequencing diverse genomes is needed.

Here we report the long-read genome assembly of *Momordica charantia*, currently the most complete assembly among publicly available genomes in Cucurbitaceae. With population re-sequencing, we also investigated the genetic structure, demographic history, patterns of selection, as well as genomic regions associated with important fruit traits.

## Results

### Genome architecture

We used PacBio contigs and two linkage maps to construct the chromosome-level genome of *M. charantia.* In total, 2,366,274 subreads with 10,725 bp read on average, equivalent to 25.3 Gb, were acquired. The genome was assembled into 302.99 Mb in 221 contigs with 96.4% BUSCO completeness. For chromosome map development, two independent linkage maps were reconstructed using previously analyzed RAD-seq (Restriction-site Associated DNA Sequence) data of two F_2_ populations (Urasaki *et al.* 2017; Cui *et al.* 2018). After imputing missing marker genotypes in F_2_ populations, we identified 12 linkage groups from the OHB61-5 x OHB95-1A cross (Urasaki *et al.* 2017) and 10 linkage groups from the K44 x Dali-11 cross (Cui *et al.* 2018). The final set of 11 chromosomes were identified by comparison between the two linkage maps. By mapping *de novo* assembly against this chromosome map, 96.27% of the sequences (291.7Mb) can be anchored in chromosomes. Comparing among all published Cucurbitaceae genomes (including a recent Illumina-based *M. charantia* assembly of the Dali-11 accession, European nucleotide archive PRJEB24032), our assembly has the highest contig N50 (close to 10 Mb) as well as highest proportion of sequences placed on chromosomes (Table 1). Comparison between our long-read and the recent short-read assemblies revealed some incongruity near the centromeric regions, and much of the centromeric regions in the long-read assembly is absent from the short-read assembly (Fig. 1 and Supplementary Fig. 1).

**Table 1.** Comparison of genome assemblies in Cucurbitaceae.

|  | *Cucumis sativus* | *Citrullus lanatus* | *Citrullus lanatus* (charleston gray) | *Cucurbita argyrosperma* | *Cucurbita pepo* | *Lagenaria siceraria* | *Cucumis melo* | *Momordica charantia* (2017 assembly) | *Momordica charantia* (Dali-11 assembly) | *Momordica charantia* (this study) |
| --- | --- | --- | --- | --- | --- | --- | --- | --- | --- | --- |
| assembly size (Mb) | 243 | 353 | 396 | 229 | 263 | 313.4 | 375 | 285 | 294 | 303 |
| Contig N50 (kb) | 19.8 | 26.4 | 36.7 | 463.4 | 110.0 | 28.3 | 60.8 | 21.9 | 62.6 | 9,898.0 |
| scaffold N50 (kb) | 1,140 | 2,380 | 7,471 | 620 | 1,750 | 8,701 | 4,680 | 1,101 | 611 | 9,898 |
| sequence on chromosome (Mb) | 176.9 | 330 | 378.7 | 0 | 214.1 | 297.6 | 337.5 | 0 | 251 | 291.7 |
| sequence on chromosome / assembly size (%) | 72.80 | 93.48 | 95.63 | 0.00 | 81.41 | 94.96 | 90.00 | 0.00 | 85.37 | 96.27 |

**Figure 1.**
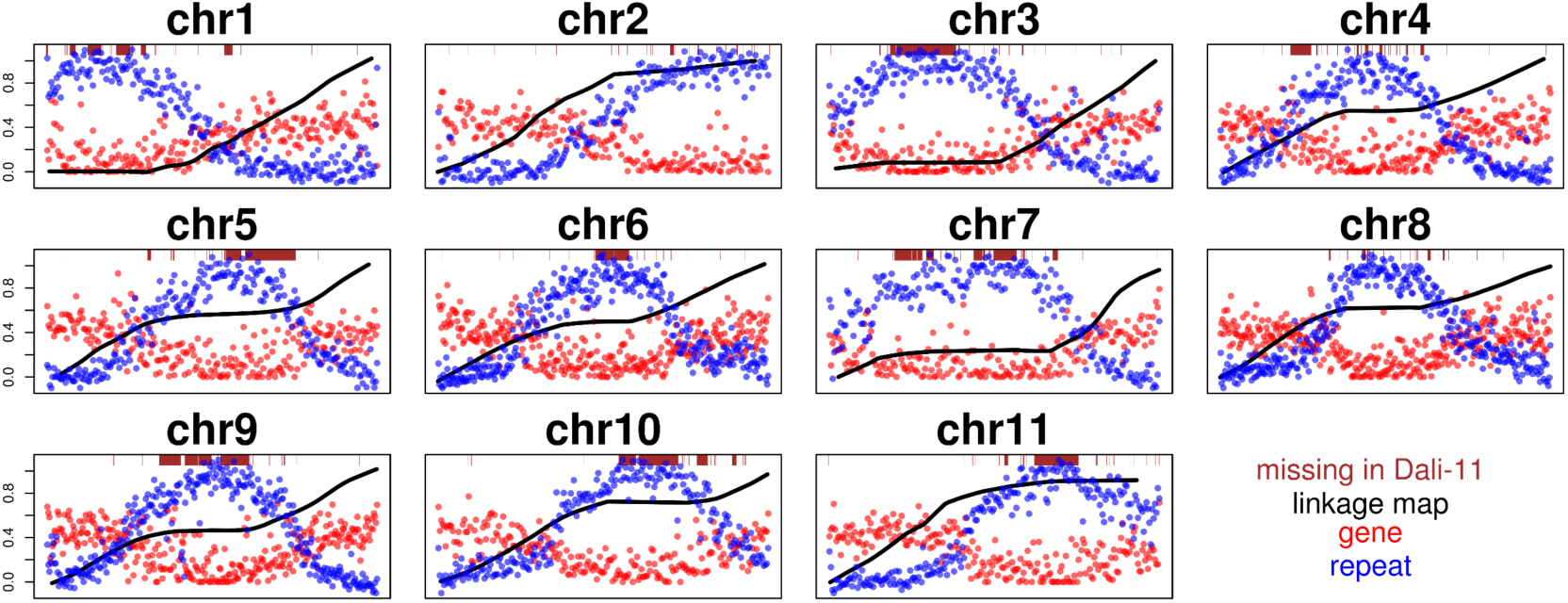
Features of the Goya v2 assembly. Shown are the genome-wide distribution of gene and repeat density, linkage map, and missing region in the Dali-11 genome. Centromeric regions inferred from the linkage map are consistent with the repeat distribution and regions absent in the Dali-11 genome.

We identified 159Mb repetitive elements (RE), representing 52.52% of the genome (Supplementary Table 1). Using the same repeat annotation pipeline, we found the repeat coverage in our assembly is higher than the Dali-11 *M. charantia* assembly (45.43%) and the other Cucurbitaceae species, demonstrating the better assembly of repetitive regions. Compared to Dali-11, long-terminal repeats (LTR), representing about 24% of the genome and 46% of all REs, are largely responsible for the higher RE proportion in our assembly (Supplementary Table 2). *Gypsy* and *Copia* subfamilies constitute most of the LTRs (25.6% and 15.8% of REs).

We further plotted the genome-wide distribution of each type of repeats. LTR, DNA transposons, and unknown repeats are enriched near the centromeric regions, representing the major improvement of long-read over the short-read assembly (Supplementary Fig. 2a-c). For other repeat categories, Short interspersed nuclear elements (SINEs) and simple-repeats have similar distribution patterns to genes (Supplementary Fig. 2d, e), and rRNAs had six unique clusters in the genome (Supplementary Fig. 2f). Interestingly, while LTRs are concentrated near the centromere, DNA transposons and long interspersed nuclear elements (LINEs) have a more peri-centromeric distribution pattern (Supplementary Fig. 2 b, g).

Our well-assembled genome also allows the synteny comparison between *M. charantia* and six other Cucurbitaceae species (Fig. 2). Between the bitter gourd and other cucurbit genomes, in general there is not a one-to-one relationship in chromosomes, indicating that these Cucurbitaceae species do not have similar karyotype to the bitter gourd. It is worth noting that in our assembly, the repeat-rich peri-centromeric regions often find little match on other genomes, again demonstrating we have assembled regions that were previously difficult for short-read genomes. Highly conserved synteny between bitter gourd and melon (*Cucumis melo*) could be observed in two pairs of chromosomes (chr1 and chr8, chr3 and chr12). Particularly, according to dotplots (Supplementary Fig. 3), more than 8Mb of euchromatic region in the end of bitter gourd chr1 showed conserved synteny with chromosomes in all analyzed Cucurbitaceae plants, while inversions were sometimes observed. In *Cucurbita maxima* and *Cucurbita moschata*, bitter gourd chr1 is in syntenic to their chr3 and chr7, implying genome duplications in *Cucurbita* species.

**Figure 2.**
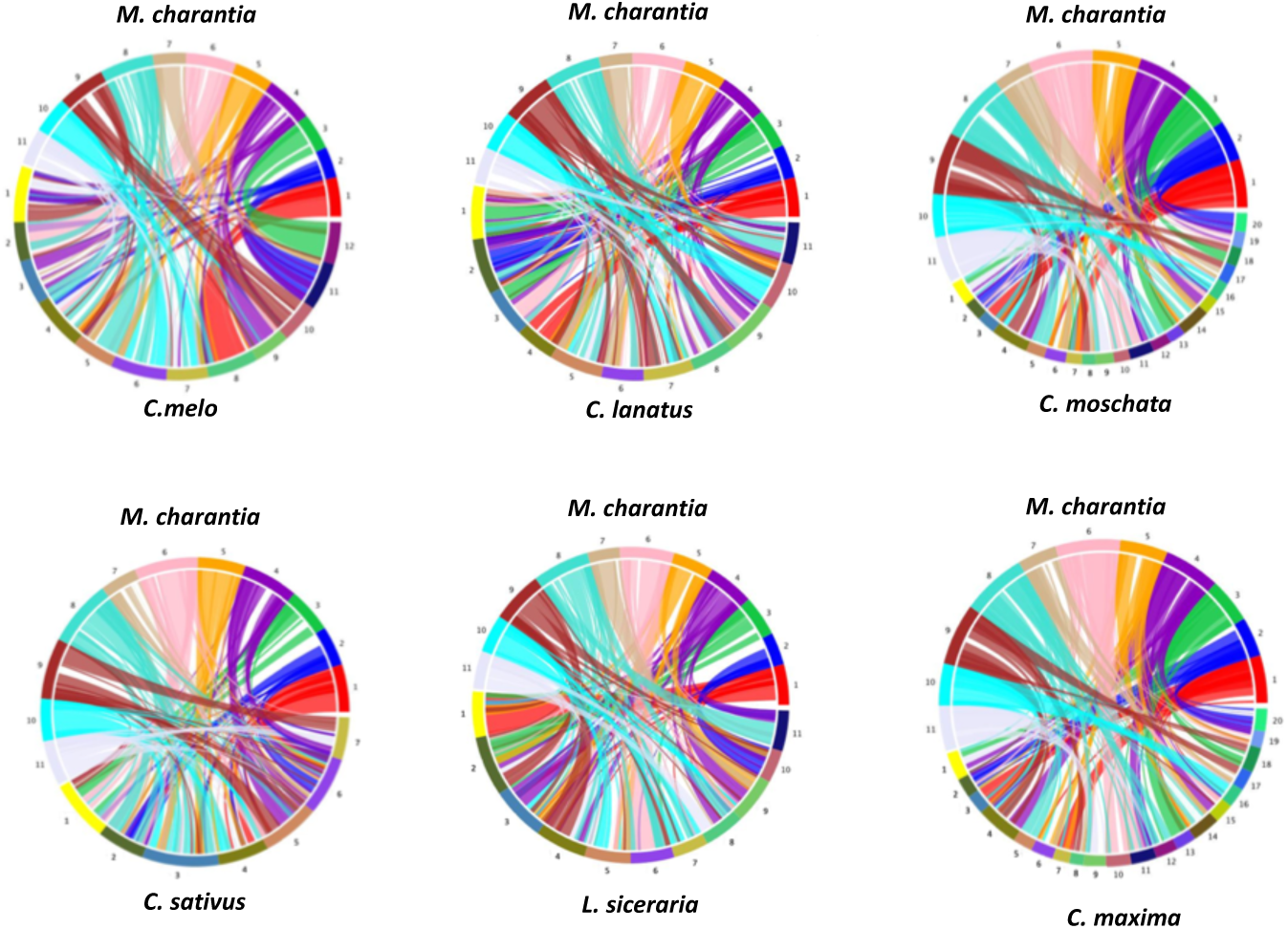
Synteny comparison between our *M. charantia* assembly and six other Cucurbitaceae species, *Cucumis melo*, *Citrullus lanatus*, *Cucurbita moschata*, *Cucumis sativus*, *Lagenaria siceraria*, and *Cucurbita maxima*.

### Demographic history

We sampled 42 cultivars, 18 wild accessions, and an outgroup (*Momordica cochinchinensis*). Population genetic analyses from ADMIXTURE (Fig. 3a), neighbor-joining tree (Fig. 3b), and principal component analysis (PCA) (Fig. 3c) consistently identified four genetic groups, including two cultivar groups from South Asia (SA) and Southeast Asia (SEA) as well as wild genetic groups from Taiwan (TAI) and Thailand (THAI). These methods give largely consistent results, with ADMIXTURE K=2 first separated wild and cultivar groups, follow by K=3 separating the two cultivar groups. Under K=5, the two wild groups as well as a small subgroup, Bangladesh within the SA group, were further separated. Correspondingly, the ADMIXTURE models had lower cross-validation errors under K=3 or 5 (Supplementary Fig. 4). We did not observe any admixed individual between TAI and THAI groups probably due to the dis-continuous spatial sampling, and the cultivar-wild admixture accessions (admix-wild) consistently possess introgressions from wild groups of the same geographic area.

**Figure 3.**
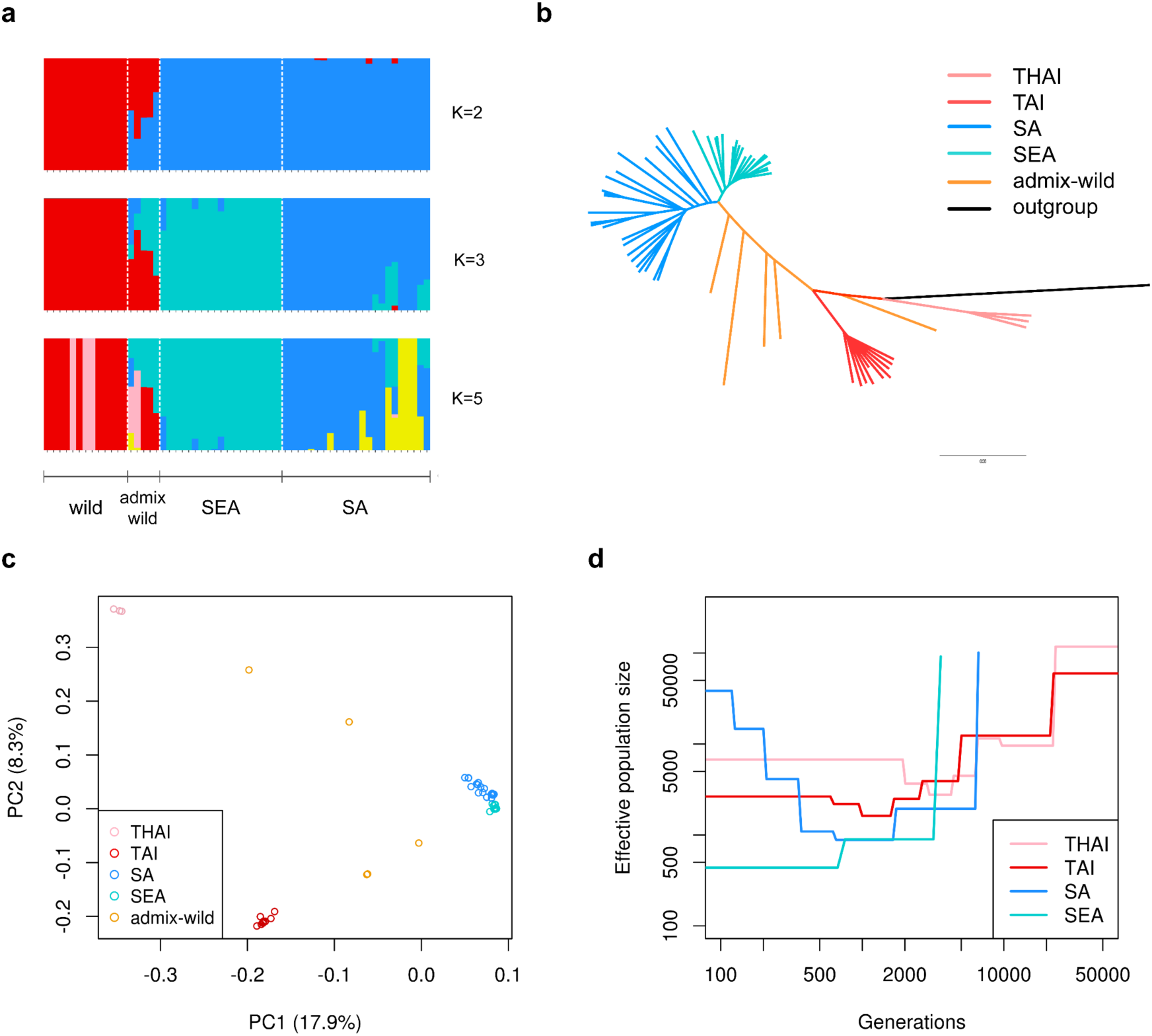
Population structure and demographic history of *Momordica charantia*. Shown are the (a) population structure, (b) neighbor-joining tree, (c) principal component analysis, and (d) demographic history of different wild (THAI and TAI) and cultivar (SA and SEA) groups.

As expected, the wild group has most rapid decay of linkage disequilibrium (LD) among the un-admixed groups, reaching low LD (*r*^2^=0.25) at about 10 kb. The pattern is seconded by SA (at about 670 kb) and SEA cultivars (about 1 Mb) (Supplementary Fig. 5). Consistent with the patterns of LD decay, the wild group has highest mean pairwise nucleotide diversity and heterozygous sites among all three groups while SEA has the lowest (Supplementary Fig. 6 and 7), suggesting the SEA cultivars represent a more recent split from the SA cultivars. Finally, the admix-wild group consisting of admixed accessions between wild and cultivars contains the highest variation and heterozygosity, consistent with their hybrid origin.

We used SMC++ (Terhorst *et al.* 2017) to infer the divergence time among these groups. The cultivars diverged from the wild groups at about 6100 years ago, and the divergence between SA and SEA cultivars happened much more recently, roughly 800 years ago (Fig. 3d and Supplementary Fig. 8).

### Genetics and signatures of selection

We employed four methods to investigate signatures of selection during the domestication process: the composite likelihood ratio test (CLR) within the cultivars and the fixation index (*F*_*ST*_), reduction of diversity (ROD), and cross-population composite likelihood ratio test (XP-CLR) between wild accessions and cultivars (Fig. 4). In general, we did not observe strong agreements among these methods, except one region in chromosome 7 associated with fruit color (below). From each method we further chose the top 1% regions and investigated the enrichment of gene ontology (GO) functional groups. In general, GO terms associated with metabolic processes are enriched in the genomic regions with top scores of these selection tests, suggesting the wild and cultivar groups are highly differentiated in metabolism-related traits (Supplementary Fig. 9).

**Figure 4.**
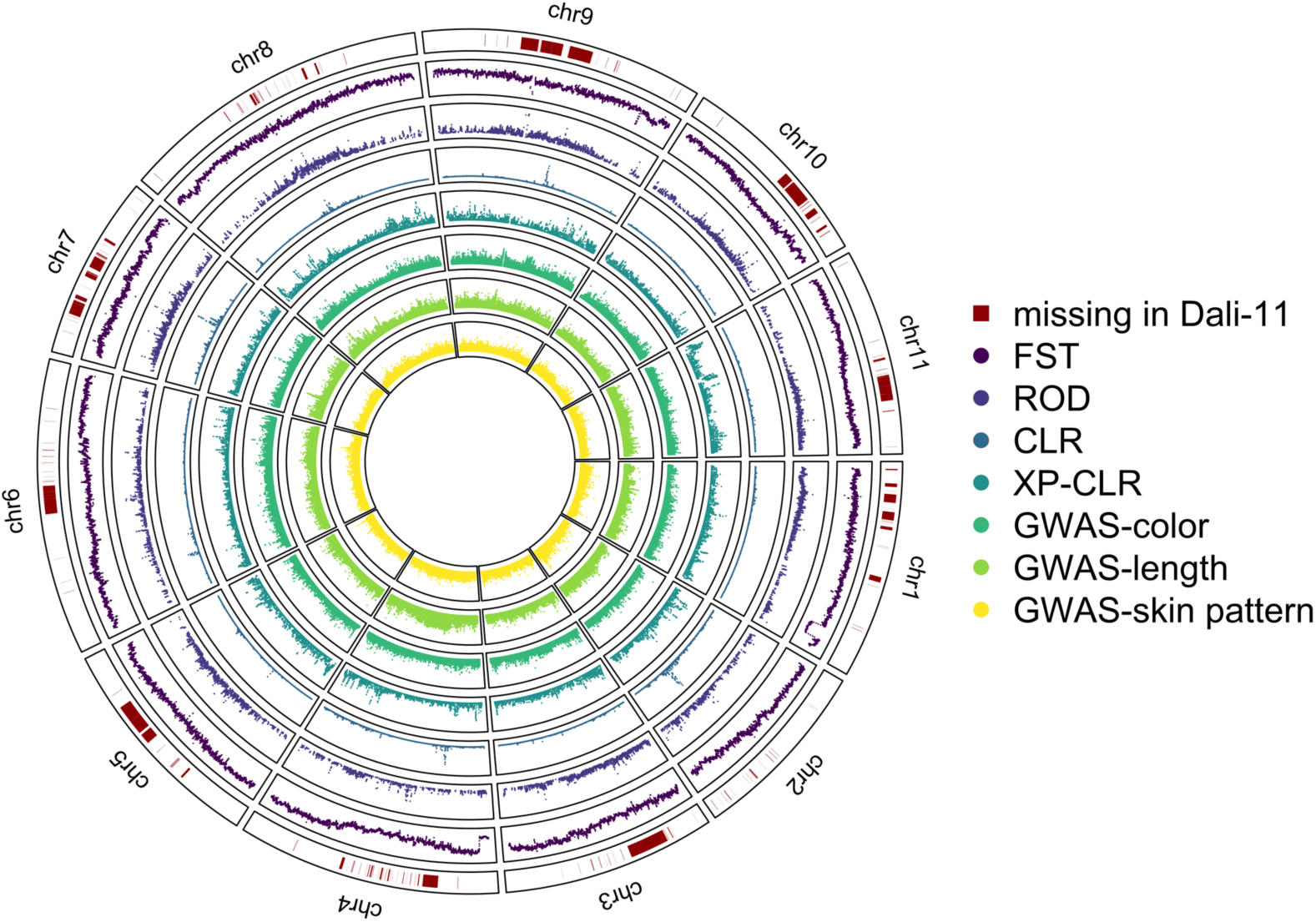
Genome-wide plot of missing regions in the Dali-11 genome, *F*_*ST*_, reduction of diversity (ROD), composite likelihood ratio (CLR), cross-population composite likelihood ratio (XP-CLR), and genome-wide association study (GWAS) results of fruit color, length, and skin pattern.

Given the high divergence between wild and cultivar groups, the baseline *F*_*ST*_ is too high to show obvious peaks. On the other hand, we observed two regions with exceptionally low *F*_*ST*_, one near the end of chromosome 1 and the other at the beginning of chromosome 4 (Fig. 4), suggesting strong selection preventing the divergence between wild and cultivar groups. Interestingly, the end of chromosome 1 harbors a locus for gynoecy, affecting the ratio of male and female flowers in this monoecious species. The locus was identified in a cross between Japanese accessions OHB61-5 and OHB95-1A (Matsumura *et al.* 2014), and our re-analyses identified two closely linked quantitative trait loci (QTL) in this region, where the QTL with larger effect (with LOD score > 30) completely overlapped this low-*F*_*ST*_ region (Fig. 5a). QTL in the same region were also identified in another cross between Chinese accessions Dali-11 and K44 (Cui *et al.* 2018). While increasing the proportion of female flowers is the focus of continuous breeding efforts in the accession level, in the population level this locus is expected to be under negative frequency dependent selection as either the fixation or loss of female-biased allele results in overall lower fitness. As expected from balancing selection, levels of polymorphism in this low-*F*_*ST*_ region is high in both wild and cultivar groups (Fig. 5a). Phylogenetic reconstruction revealed at least three distinct allelic groups in this region, and both wild and cultivar accession groups possess at least two of the three allelic groups (Fig. 5b). The balanced distribution of these highly differentiated allelic groups among populations is also consistent with patterns of balancing selection. While most studies reported how domestication efforts left strong signatures of artificial selection in the genome, here we report a likely case where human efforts could not overcome balancing selection in nature.

**Figure 5.**
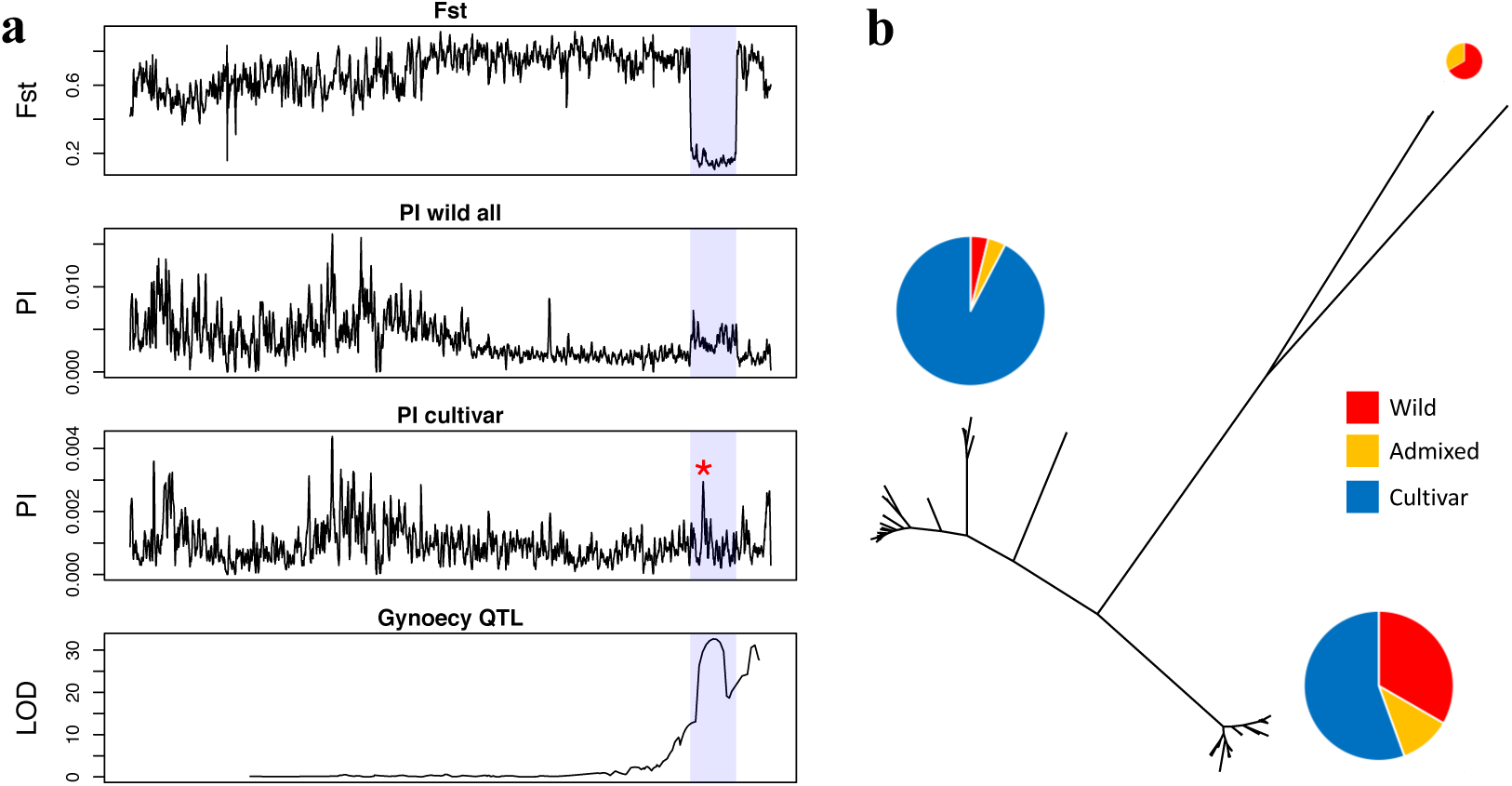
The region with low differentiation and high polymorphism between wild and cultivar groups in chromosome 1. (a) This region in chromosome 1 is labeled in the shaded area, showing low *F*_*ST*_, high PI, and the co-localization with the gynoecy QTL. The red asterisk labels the region for phylogenetic tree reconstruction in (b). (b) Phylogenetic tree of the high diversity region, showing at least three highly balanced allelic groups. The pie charts represent proportion of the wild, wild-cultivar admixed, and cultivar accessions (assigned from genomewide SNPs) containing these alleles.

To investigate the genetic architecture underlying traits associated with domestication, genome-wide association study (GWAS) was performed on skin pattern, fruit color, and fruit length (Figure 4 and 6, Supplementary Table 3). From wild to SA to SEA groups, in general the fruits became less spiny, lighter in color, and larger. Fruit color ranges from white to dark green, with the wild accessions having green and the cultivars having lighter but sometimes darker fruits (Fig. 6a). We found strong traces of selection for Peak3 in chromosome 7 (Fig. 6a and 7a) where the SA group is enriched for the allele associated with darker fruits. Under this peak are two candidate genes (*Y3093_ARATH*) with proteins homologous to the chloroplastic gene AT3G60930 in Arabidopsis. While many accessions, especially in SEA, have lighter fruit color, it is yet unclear whether the change of fruit color resulted from selection or just the consequence of genetic drift, and the strong traces of selection for the dark-fruit allele in Peak3 might be associated with other functional traits instead of color itself.

**Figure 6.**
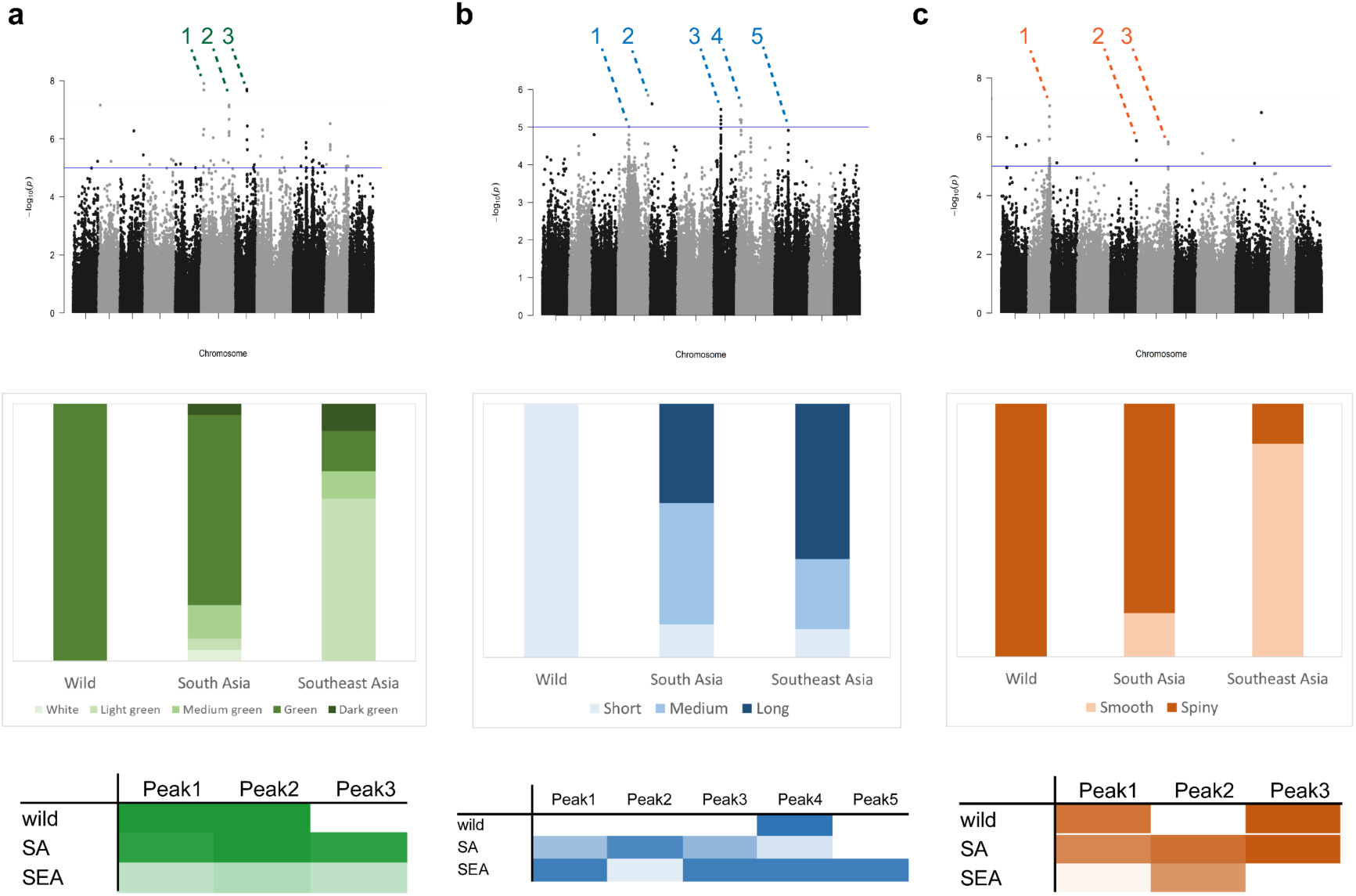
GWAS peaks and allelic effects for (a) fruit color, (b) fruit length, and (c) fruit skin pattern. The top panels are GWAS Manhattan plots with peaks labeled. The middle panels show trait distributions among genetic groups. The bottom panels show allelic effects of GWAS peaks and their allele frequency distribution: Darker colors denote higher frequency of alleles increasing trait values. GWAS peaks were chosen based on their scores as well as evidences supported by linked SNPs.

**Figure 7.**
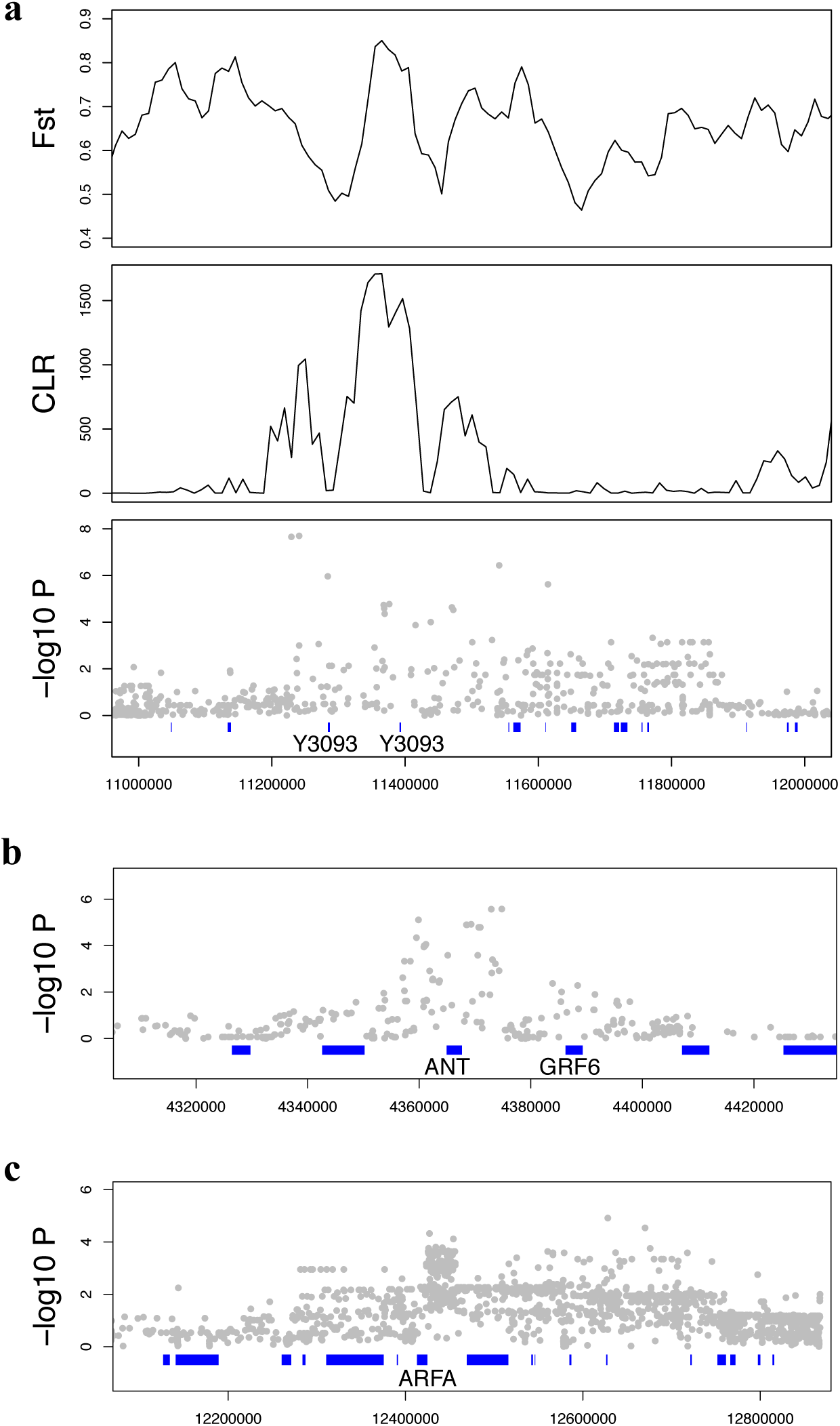
Candidate genes under genome-wide association study (GWAS) peaks and their signatures of selection. (a) There are two copies of the chloroplast gene, *Y3093*, under the peak of *F*_*ST*_, composite likelihood ratio test (CLR), and the GWAS for fruit color. Also shown are GWAS peak and candidate genes of fruit length in (b) chromosome 8 and (c) chromosome 9.

For fruit size, while the size-increasing allele has highest frequency in the SA group for Peak2, in other GWAS peaks the SEA group has more size-increasing alleles, corresponding to its generally larger fruit size (Fig. 6b). This is consistent with the classic model of polygenic selection where parallel selections for larger fruits in SEA and SA resulted in the allele frequency shifts in independent loci. Our observation that SEA and SA cultivars harbor size-increasing alleles in independent loci also echoes previous observations that heterosis exists in inter-group crosses in this species. While further studies are required to validate these GWAS peaks, we identified several promising candidate genes (Fig. 7b). For example, under Peak4 in chromosome 8 we found *ANT*, an ethylene-responsive transcription factor required for the development of female gametophyte, ovule integument, and gynoecium marginal tissues (Klucher *et al.* 1996; Liu *et al.* 2000; Mizukami and Fischer 2000), and *GRF6*, the Growth-regulating factor with regulatory roles in gibberellin-induced growth (Tanaka *et al.* 2008). Under Peak5 in chromosome 9 we found *ARFA*, an Auxin response factor that promotes flowering, stamen development, floral organ abscission and fruit dehiscence (Hagen and Guilfoyle 2002; Ellis *et al.* 2005). Under one of the highest peaks (peak 3 in chromosome 7, Supplementary Fig. 10), we found genes associated with signal transduction (CRK3), fatty acid desaturation (ADS3), malate transportation (ALMT10), protein chaperon (CIA30), and flowering time (FLK). Interestingly, the genomic region containing this largest peak in our assembly is absent from the Illumina-based Dali-11 assembly (Supplementary Fig. 10).

## Discussion

In this study, we used long-read technology to assemble the genome of bitter gourd, currently the most complete assembly among all public Cucurbitaceae genomes. Our assembly contains many repeat-rich regions not assembled by the recent Dali-11 genome (European nucleotide archive PRJEB24032). While we are yet to pinpoint the candidate genes directly associated with plant size, one of the highest GWAS peaks for fruit size resides within the region missing from the short-read assembly (Supplementary Fig. 10), demonstrating the importance of a well-assembled long-read genome.

Unlike “classic examples” of domestication such as the loss of branching in maize and the loss of shattering in rice, the direction of selection in bitter gourd is not ubiquitous: Consumers from different cultures have their own preferences. South Asians like highly bitter fruit with smaller size (although still larger than wild accessions) and spiny and dark green features. Southeast Asians like less bitter fruit with light green (or white) and smooth skin pattern (Dhillon *et al.* 2016). Considerable phenotypic variations therefore exist for *M. charantia* cultivars. The situation in bitter gourd may be more akin to the high diversity of local maize landraces in Central America, rather than the commercial elite maize cultivars produced globally. Under such situation, it is therefore conceivable that it would be difficult to identify classic signatures of selective sweep and strong Mendelian genes, given that the most obvious target of selection is fruit size, likely a polygenic trait. The situation may be further complicated by that some cultures value and cultivate the wild accessions for their bitterness. Therefore, the process of selection may be slower in bitter gourd, with frequent introgressions between wild and cultivar groups, preventing the strong and rapid fixation of underlying genes.

Our analyses showed that the SA group first diverged from wild progenitors at about 6000 years ago, followed by the splitting of the SEA group about 800 years ago. Consistent with the aforementioned cultural preference, the SEA group is enriched for alleles conferring lighter skin color, larger fruits, and smoother fruit surface compared to the SA group. We also performed genomic tests of signatures of selection for the SEA group only (data not shown), but no strong pattern was observed either. This is likely due to the underlying genetic architecture of selected traits being polygenic. Interestingly, we found one region with very low divergence between wild and cultivar groups likely colocalizing with the locus conferring gynoecy. Despite being a continuous focus of breeding efforts (Matsumura *et al.* 2014), the need to keep balanced sex ratio in nature may prevent the divergence between wild and cultivar groups in this region. Taken together, our investigations showed that the bitter gourd may provide a valuable “non-classic” model of domestication, where the intermittent weak selection and polygenic genetic architecture precludes the identification of strong candidate genes, and the directional artificial selection for gynoecy cannot overwhelm the forces of balancing selection in nature.

## Materials and Methods

### Genome assembly

High molecular weight genomic DNA was extracted from the leaves of *M. charantia* OHB3-1 accession following the protocol provided from Pacific Biosciences with modification. Briefly, genomic DNA was extracted from the leaf tissue using Carlson Lysis buffer containing CTAB and precipitated by ethanol after chloroform/isoamyl alcohol extraction. RNase- and proteinase-treated genomic DNA was purified using Genomic-tip (Qiagen). SMART library was prepared from high molecular weight genomic DNA (>50kbp) and applied to sequencing by PacBio Sequel.

Subreads from PacBio sequencing were corrected and assembled using Canu 1.7 with default settings for PacBio. The obtained contigs were polished by pilon 1.23 using paired-end Illumina HiSeq2500 reads (250b x2) from the same genomic DNA.

Restriction-site Associated DNA Sequence (RAD-seq) (Davey and Blaxter 2010) data were also obtained from two F2 crosses: 97 F2 individuals from a cross in Japan (Urasaki *et al.* 2017) and 423 F2 individuals from a cross in China (Cui *et al.* 2018). In order to solve the low coverage and high missing-data problem in RAD-seq data, we employed a window-based method to define marker genotypes (Lee *et al.* 2017). Briefly, the genome was cut into 100-kb windows, and the parental genotype of each F2 individual within each window was called based on the proportion of parental reads within the window: If the maternal allele proportion is higher than 80%, we set it to maternal homozygous; if the proportion is lower than 20%, we set it to paternal homozygous; if the proportion is between 20% and 80%, we defined the genotype as heterozygous. We used the same filter conditions in both crosses. SNPs with allele depth (AD) <3 and maternal allele proportion >= 95% or <=5% across all samples were excluded. For 100-kb windows used for linkage map construction, if the depth of a sample in a window is lower than 5, we called it missing, and a window was excluded if the proportion of missing individuals is higher than 60%.

MSTmap (Wu *et al.* 2008) was used for constructing linkage maps. After running MSTmap, we filtered possible genotyping errors: In an individual, if two recombinant events happened within 10 cM, we set the genotypes of markers between the two breakpoints to missing. We also imputed the missing markers: If the upstream and downstream of a missing marker has the same allele, and this missing marker is within 10 cM of either two closest markers, we imputed it with the closest marker. Consequently, we identified 12 linkage groups from the Japanese cross (Urasaki *et al.* 2017) and 10 linkage groups from the Chinese cross (Cui *et al.* 2018). The final set of 11 linkage groups were identified by comparison between the two linkage maps. Finally, we redid MSTmap for each linkage group separately and generated the final linkage map. ALLMAPS (Tang *et al.* 2015) was used to combine the two linkage maps and produce the chromosome-level assembly. Scaffolds smaller than 10 kb were excluded from the construction. We set the weight of Japan linkage map to 1.5 and Chinese map to 1 since our genome strain was genetically closer to parents in the Japan cross. MUMmer (Kurtz *et al.* 2004) was used to align the Dali-11 (European nucleotide archive PRJEB24032) assembly and our assembly.

Synteny blocks were identified between the genomes of bitter gourd and other Cucurbitaceae species. Sequences of pseudomolecules and GFF files for the predicted genes in *Cucumis melo* (Argyris *et al.* 2015), *Cucumis sativus* (Li *et al.* 2019), *Citrullus lanatus* (Guo *et al.* 2013), *Cucurbita maxima*, *Cucurbita moschata* (Sun *et al.* 2017), and *Lagenaria siceraria* (Wu *et al.* 2017) were applied to the analysis by SyMAP 4.2 (Soderlund *et al.* 2006; Soderlund *et al.* 2011) with default settings.

### Gene annotation

We performed repeat annotation by RepeatMasker (Tarailo-Graovac and Chen 2009) with a *de novo* repeat library constructed by RepeatModeler (Smit Afa 2008–2015) and Repbase (Bao *et al.* 2015). We used *ab-initio* gene prediction and RNA-seq data for gene annotation. RNA-seq data from three tissues, root (SRR3535149), leaf (SRR3535138), and flower (SRR3535137) were mapped to the genome by HISAT2 (Kim *et al.* 2015) and subsequently assembled and merged by StringTie (Pertea *et al.* 2015). We used TransDecoder (Haas *et al.* 2013) to predict the open reading frame (ORF) based on assembled transcripts, followed by the use of parameter “retain_blastp_hits” to validate the result using blastp (Altschul *et al.* 1990) on UniProt (Uniprot 2019) database. *Ab-initio* gene prediction by AUGUSTUS (Stanke and Morgenstern 2005) was performed with the repeat-masked genome with “-species Arabidopsis” option. The species parameters of AUGUSTUS were trained by genome mode BUSCO (Waterhouse *et al.* 2018) with eudicotyledons_odb10 database.

The *Ab-initio* predictions, RNA-seq alignments, and ORF predictions were submitted to Evidencemodeler (EVM) (Haas *et al.* 2008) to identify consensus gene model. The weight of *ab-initio* and ORF prediction is 1, and RNA-seq data have a weight of 10, based on the recommendation of EVM. The gene set from EVM was sent to BUSCO for assessing the completeness with eudicotyledons_odb10 database.

The complete gene set was loaded into blast2go (Gotz *et al.* 2008) and compared with UniProtKB/Swiss-prot (Uniprot 2019) database using local blastx. Blast E-expectation value (E-value) cutoff was set to 0.001 and word size to 6. Moreover, we mapped the genes annotated by blastx to the GO database. The mapped GO terms were further evaluated by GO evidence codes, which indicated the experimental and computational evidence of GO terms. GO enrichment analysis of genomic regions with signatures of selection was implemented with Fisher’s Exact Test (Jowett 1956).

### Plant materials and population genetics analyses

A total of 60 *Momordica charantia* accessions were used for population genomics analyses, with 45 cultivars and 12 wild accessions from the World Vegetable Center and 3 wild ones collected in Taiwan. The outgroup, *Momordica cochinchinensis*, were obtained from a horticulture market in Taiwan, and its species identity was subsequently validated with chloroplast *MaturaseK* gene (*MatK*) markers (Supplementary Table 4) (Ka *et al.* 1995). The phenotypic data were received from the World Vegetable Center in Thailand. All of them are categorical and graded data (Supplementary Table 3). The estimation method of the phenotypes had been reported in a previous study (Dhillon *et al.* 2016).

The genomic DNA was extracted from leaves using DNeasy Plant Mini Kit (Qiagen) with 100 mg of leaf tissue, and DNA quality and concentration were estimated with gel electrophoresis and Qubit. NEBNext Ultra II DNA Library Prep Kit was used to construct the illumina library. Library size was selected to be around 500 bp. The libraries were sequenced with 150 bp paired-end using Illumina HiSeq X-ten.

Reads were trimmed base on sequence quality by SolexaQA (Cox *et al.* 2010), and adaptor sequences were removed by cutadapt (Martin 2011). Reads were mapped to the reference genome by BWA 0.7.15 (Li and Durbin 2010). The duplicated reads produced by PCR were marked with Picard Tools (http://broadinstitute.github.io/picard). SNP genotypes were called following GATK3.7 (Mckenna *et al.* 2010) best practice. Variant sites were then filtered with vcftools (Danecek *et al.* 2011) by keeping the bi-allelic SNP sites only, QUAL > 30, missing rate < 10%, and minor allele frequency (MAF) > 1%. Sites with depth among all samples lower or higher than 3 standard deviations of genome-wide average were filtered out. For genome-wide association study (GWAS) analysis, the alleles with MAF < 10% were excluded.

PLINK (Purcell *et al.* 2007) was used to perform SNP linkage disequilibrium (LD) pruning in 50-kb windows, 5-kb between each step, and *r*^2^ threshold of 0.5. The neighbor-joining tree was reconstructed with TASSEL (Bradbury *et al.* 2007) and visualized with FigTree v.1.4.3 (http://tree.bio.ed.ac.uk/software/figtree/). Principal component analysis (PCA) was performed by PLINK (Purcell *et al.* 2007) with default settings. Ancestral proportion analysis was performed by ADMIXTURE (Alexander *et al.* 2009) with “--cv” to estimate cross-validation error of each K value, and the admixture Q matrix was plotted by R package pophelper (Francis 2017).

LD decay was calculated and plotted with PopLDdecay (Zhang *et al.* 2019). We removed the admixed individuals identified by ADMIXTURE before LD estimation within each genetic group. Nucleotide diversity was calculated in 50-kb windows with 10-kb steps by vcftools. Heterozygosity of each individual was counted by vcftools with “--het” option.

We used SMC++ (Terhorst *et al.* 2017) to estimate the demographic history of *M. charantia*. SMC++ had two advantages: 1) It required only unphased genomes, which was suitable for non-model organisms. 2) Multiple samples could be included in the analysis for constructing the recent past. The admixed individuals in each group were excluded before analyses. Historical population sizes of four genetic groups, THAI, TAI, SA, and SEA were separately estimated with the “estimate” option, and their pairwise divergence times were estimated by “split” option. After summarizing the mutation rates frequently used for eudicots, the mutation rate per generation was set as 2×10^−8^.

The wild group we used in selection models was the Taiwan wild group since it was genetically closer to the cultivars. The fixation index (*F*_*ST*_) between wild and cultivar populations was calculated in 50-kb windows with 10-kb step size by vcftools. Reduction of diversity (ROD) was calculated in 50-kb window with 10-kb step size between the wild and cultivar populations. The formula was: *log*_*10*_ (*π*_*wild*_/*π*_*cultivar*_). Composite likelihood ratio (CLR) (Zhu and Bustamante 2005) was performed within cultivars by SweeD (Pavlidis *et al.* 2013). Each chromosome was separated into 2,000 bins. Cross-population composite likelihood ratio (XP-CLR) (Chen *et al.* 2010) was estimated between wild and cultivar populations in 50-kb windows with 10-kb step size by xpclr software (https://github.com/hardingnj/xpclr). The genetic distance between markers was calculated with Japanese linkage map by interpolation. Genome-wide association study (GWAS) was performed with the general linear model in TASSEL (Bradbury *et al.* 2007). The manhattan plots were plotted by R package qqman (Turner 2014)

## Acknowledgements

The authors thank Chia-Yu Chen, Pei-Min Yeh, Jo-Wei Hsieh, and Jo-Yi Yen for assistance. We are grateful to the support from National Taiwan University’s Computer and Information Networking Center for high-performance computing facilities and College of Life Science Technology Commons for molecular biology facilities. This work was supported by Taiwan Ministry of Science and Technology grant number 107-2636-B-002-004 and 108-2636-B-002-004, JSPS KAKENHI Grant Number JP17K07601, and long-term strategic donors to the World Vegetable Center: Republic of China (Taiwan), UK aid from the UK government, United States Agency for International Development (USAID), Australian Centre for International Agricultural Research (ACIAR), Germany, Thailand, Philippines, Korea, and Japan..

## Data availability

The assembled genome was available under DNA Data Bank of Japan, accession number. The PacBio reads were submitted under DNA Data Bank of Japan, accession number DRR194981. The Illumina reads of the OHB3-1 genome strain was submitted under DNA Data Bank of Japan, accession number DRR194976. Population re-sequencing Illumina reads were submitted under NCBI BioProject PRJNA578358.

## References

Alexander, D. H., J. Novembre, and K. Lange, 2009 Fast model-based estimation of ancestry in unrelated individuals. Genome Res 19: 1655–1664.

Altschul, S. F., W. Gish, W. Miller, E. W. Myers, and D. J. Lipman, 1990 Basic local alignment search tool. J Mol Biol 215: 403–410.

Argyris, J. M., A. Ruiz-Herrera, P. Madriz-Masis, W. Sanseverino, J. Morata et al., 2015 Use of targeted SNP selection for an improved anchoring of the melon (Cucumis melo L.) scaffold genome assembly. Bmc Genomics 16: 4.

Bao, W., K. K. Kojima, and O. Kohany, 2015 Repbase Update, a database of repetitive elements in eukaryotic genomes. Mob DNA 6: 11.

Bradbury, P. J., Z. Zhang, D. E. Kroon, T. M. Casstevens, Y. Ramdoss et al., 2007 TASSEL: software for association mapping of complex traits in diverse samples. Bioinformatics 23: 2633–2635.

Chen, H., N. Patterson, and D. Reich, 2010 Population differentiation as a test for selective sweeps. Genome Research 20: 393–402.

Cox, M. P., D. A. Peterson, and P. J. Biggs, 2010 SolexaQA: At-a-glance quality assessment of Illumina second-generation sequencing data. Bmc Bioinformatics 11.

Cui, J., S. Luo, Y. Niu, R. Huang, Q. Wen et al., 2018 A RAD-Based Genetic Map for Anchoring Scaffold Sequences and Identifying QTLs in Bitter Gourd (Momordica charantia). Frontiers in Plant Science 9: 477.

Danecek, P., A. Auton, G. Abecasis, C. A. Albers, E. Banks et al., 2011 The variant call format and VCFtools. Bioinformatics 27: 2156–2158.

Davey, J. L., and M. W. Blaxter, 2010 RADSeq: next-generation population genetics. Briefings in Functional Genomics 9: 416–423.

Dhillon, N. P. S., S. Sanguansil, R. Schafleitner, Y.-W. Wang, and J. D. McCreight, 2016 Diversity Among a Wide Asian Collection of Bitter Gourd Landraces and their Genetic Relationships with Commercial Hybrid Cultivars. Journal of the American Society for Horticultural Science 141: 475–484.

Ellis, C. M., P. Nagpal, J. C. Young, G. Hagen, T. J. Guilfoyle et al., 2005 AUXIN RESPONSE FACTOR1 and AUXIN RESPONSE FACTOR2 regulate senescence and floral organ abscission in Arabidopsis thaliana. Development 132: 4563–4574.

Francis, R. M., 2017 pophelper: an R package and web app to analyse and visualize population structure. Mol Ecol Resour 17: 27–32.

Gaikwad, A. B., T. K. Behera, A. K. Singh, D. Chandel, J. L. Karihaloo et al., 2008 Amplified fragment length polymorphism analysis provides strategies for improvement of bitter gourd (Momordica charantia L.). Hortscience 43: 127–133.

Gotz, S., J. M. Garcia-Gomez, J. Terol, T. D. Williams, S. H. Nagaraj et al., 2008 High-throughput functional annotation and data mining with the Blast2GO suite. Nucleic Acids Research 36: 3420–3435.

Guo, S. G., J. G. Zhang, H. H. Sun, J. Salse, W. J. Lucas et al., 2013 The draft genome of watermelon (Citrullus lanatus) and resequencing of 20 diverse accessions. Nature Genetics 45: 51–58.

Haas, B. J., A. Papanicolaou, M. Yassour, M. Grabherr, P. D. Blood et al., 2013 De novo transcript sequence reconstruction from RNA-seq using the Trinity platform for reference generation and analysis. Nat Protoc 8: 1494–1512.

Haas, B. J., S. L. Salzberg, W. Zhu, M. Pertea, J. E. Allen et al., 2008 Automated eukaryotic gene structure annotation using EVidenceModeler and the program to assemble spliced alignments. Genome Biology 9.

Hagen, G., and T. Guilfoyle, 2002 Auxin-responsive gene expression: genes, promoters and regulatory factors. Plant Molecular Biology 49: 373–385.

Jowett, G. H., 1956 Statistical-Methods for Research Workers - Fisher,Ra. The Royal Statistical Society Series C-Applied Statistics 5: 68–70.

Ka, O., Y. Endo, J. Yokoyama, and N. Murakami, 1995 Useful primer designs to amplify DNA fragments of the plastid gene.

Kim, D., B. Langmead, and S. L. Salzberg, 2015 HISAT: a fast spliced aligner with low memory requirements. Nat Methods 12: 357–360.

Klucher, K. M., H. Chow, L. Reiser, and R. L. Fischer, 1996 The AINTEGUMENTA gene of arabidopsis required for ovule and female gametophyte development is related to the floral homeotic gene APETALA2. Plant Cell 8: 137–153.

Konishi, S., T. Izawa, S. Y. Lin, K. Ebana, Y. Fukuta et al., 2006 An SNP caused loss of seed shattering during rice domestication. Science 312: 1392–1396.

Krawinkel, M. B., and G. B. Keding, 2006 Bitter gourd (Momordica Charantia): A dietary approach to hyperglycemia. Nutr Rev 64: 331–337.

Kurtz, S., A. Phillippy, A. L. Delcher, M. Smoot, M. Shumway et al., 2004 Versatile and open software for comparing large genomes. Genome Biol 5: R12.

Lee, C.-R., B. Wang, J. P. Mojica, T. Mandáková, K. V. S. K. Prasad et al., 2017 Young inversion with multiple linked QTLs under selection in a hybrid zone. Nature Ecology & Evolution 1: 0119.

Li, H., and R. Durbin, 2010 Fast and accurate long-read alignment with Burrows-Wheeler transform. Bioinformatics 26: 589–595.

Li, Q., H. Li, W. Huang, Y. Xu, Q. Zhou et al., 2019 A chromosome-scale genome assembly of cucumber (Cucumis sativus L.). GigaScience 8: giz072.

Liu, Z. C., R. G. Franks, and V. P. Klink, 2000 Regulation of gynoecium marginal tissue formation by LEUNIG and AINTEGUMENTA. Plant Cell 12: 1879–1891.

Martin, M., 2011 Cutadapt removes adapter sequences from high-throughput sequencing reads. EMBnet.journal 17: 10.

Matsumura, H., N. Miyagi, N. Taniai, M. Fukushima, K. Tarora et al., 2014 Mapping of the Gynoecy in Bitter Gourd (Momordica charantia) Using RAD-Seq Analysis. Plos One 9: e87138.

McKenna, A., M. Hanna, E. Banks, A. Sivachenko, K. Cibulskis et al., 2010 The Genome Analysis Toolkit: a MapReduce framework for analyzing next-generation DNA sequencing data. Genome Res 20: 1297–1303.

Meyer, R. S., and M. D. Purugganan, 2013 Evolution of crop species: genetics of domestication and diversification. Nature reviews genetics 14: 840–852.

Mizukami, Y., and R. L. Fischer, 2000 Plant organ size control: AINTEGUMENTA regulates growth and cell numbers during organogenesis. Proceedings of the National Academy of Sciences of the United States of America 97: 942–947.

Pavlidis, P., D. Zivkovic, A. Stamatakis, and N. Alachiotis, 2013 SweeD: likelihood-based detection of selective sweeps in thousands of genomes. Mol Biol Evol 30: 2224–2234.

Pertea, M., G. M. Pertea, C. M. Antonescu, T. C. Chang, J. T. Mendell et al., 2015 StringTie enables improved reconstruction of a transcriptome from RNA-seq reads. Nat Biotechnol 33: 290–295.

Purcell, S., B. Neale, K. Todd-Brown, L. Thomas, M. A. Ferreira et al., 2007 PLINK: a tool set for whole-genome association and population-based linkage analyses. Am J Hum Genet 81: 559–575.

Saxena, S., A. Singh, S. Archak, T. K. Behera, J. K. John et al., 2015 Development of Novel Simple Sequence Repeat Markers in Bitter Gourd (Momordica charantia L.) Through Enriched Genomic Libraries and Their Utilization in Analysis of Genetic Diversity and Cross-Species Transferability. Applied Biochemistry and Biotechnology 175: 93–118.

Smit AFA, H. R., 2008-2015 RepeatModeler Open-1.0.

Soderlund, C., M. Bomhoff, and W. M. Nelson, 2011 SyMAP v3. 4: a turnkey synteny system with application to plant genomes. Nucleic acids research 39: e68–e68.

Soderlund, C., W. Nelson, A. Shoemaker, and A. Paterson, 2006 SyMAP: A system for discovering and viewing syntenic regions of FPC maps. Genome research 16: 1159–1168.

Stanke, M., and B. Morgenstern, 2005 AUGUSTUS: a web server for gene prediction in eukaryotes that allows user-defined constraints. 33: W465–W467.

Sun, H., S. Wu, G. Zhang, C. Jiao, S. Guo et al., 2017 Karyotype stability and unbiased fractionation in the paleo-allotetraploid Cucurbita genomes. Molecular plant 10: 1293–1306.

Tan, S. P., T. C. Kha, S. E. Parks, and P. D. Roach, 2016 Bitter melon (Momordica charantia L.) bioactive composition and health benefits: A review. Food Reviews International 32: 181–202.

Tanaka, T., B. A. Antonio, S. Kikuchi, T. Matsumoto, Y. Nagamura et al., 2008 The rice annotation project database (RAP-DB): 2008 update. Nucleic Acids Research 36: D1028–D1033.

Tang, H., X. Zhang, C. Miao, J. Zhang, R. Ming et al., 2015 ALLMAPS: robust scaffold ordering based on multiple maps. Genome Biol 16: 3.

Tarailo-Graovac, M., and N. Chen, 2009 Using RepeatMasker to identify repetitive elements in genomic sequences. Curr Protoc Bioinformatics Chapter 4: Unit 4 10.

Terhorst, J., J. A. Kamm, and Y. S. Song, 2017 Robust and scalable inference of population history from hundreds of unphased whole genomes. Nat Genet 49: 303–309.

Turner, S. D., 2014 qqman: an R package for visualizing GWAS results using Q-Q and manhattan plots. bioRxiv: 005165.

UniProt, C., 2019 UniProt: a worldwide hub of protein knowledge. Nucleic Acids Res 47: D506–D515.

Urasaki, N., H. Takagi, S. Natsume, A. Uemura, N. Taniai et al., 2017 Draft genome sequence of bitter gourd (Momordica charantia), a vegetable and medicinal plant in tropical and subtropical regions. DNA Res 24: 51–58.

Wang, R.-L., A. Stec, J. Hey, L. Lukens, and J. Doebley, 1999 The limits of selection during maize domestication. Nature 398: 236–239.

Waterhouse, R. M., M. Seppey, F. A. Simao, M. Manni, P. Ioannidis et al., 2018 BUSCO Applications from Quality Assessments to Gene Prediction and Phylogenomics. Molecular Biology and Evolution 35: 543–548.

Wu, S., M. Shamimuzzaman, H. H. Sun, J. Salse, X. L. Sui et al., 2017 The bottle gourd genome provides insights into Cucurbitaceae evolution and facilitates mapping of a Papaya ring-spot virus resistance locus. Plant Journal 92: 963–975.

Wu, Y., P. R. Bhat, T. J. Close, and S. Lonardi, 2008 Efficient and Accurate Construction of Genetic Linkage Maps from the Minimum Spanning Tree of a Graph. PLoS Genetics 4: e1000212.

Zaman, M. Y., and S. S. Alam, 2009 Karyotype Diversity in Three Cultivars of Momordica charantia L. 74: 473–478.

Zhang, C., S. S. Dong, J. Y. Xu, W. M. He, and T. L. Yang, 2019 PopLDdecay: a fast and effective tool for linkage disequilibrium decay analysis based on variant call format files. Bioinformatics 35: 1786–1788.

Zhu, L., and C. D. Bustamante, 2005 A composite-likelihood approach for detecting directional selection from DNA sequence data. Genetics 170: 1411–1421.

